# Investigating the genetic architecture of eye colour in a Canadian cohort

**DOI:** 10.1101/2021.09.29.462299

**Authors:** Frida Lona-Durazo, Rohit Thakur, Erola Pairo-Castineira, Karen Funderburk, Tongwu Zhang, Michael A. Kovacs, Jiyeon Choi, Ian J. Jackson, Kevin M. Brown, Esteban J. Parra

**Affiliations:** Department of Anthropology, University of Toronto at Mississauga, Mississauga, Ontario, Canada; Laboratory of Translational Genomics, Division of Cancer Epidemiology and Genetics, National Cancer Institute, National Institutes of Health, Bethesda, Maryland, United States of America; Integrative Tumor Epidemiology Branch, Division of Cancer Epidemiology and Genetics, National Cancer Institute, National Institutes of Health, Bethesda, Maryland, United States of America; MRC Human Genetics Unit, Institute of Genetics and Cancer, University of Edinburgh, United Kingdom; Roslin Institute, University of Edinburgh, Easter Bush, Midlothian, United Kingdom

## Abstract

The main factors that determine eye colour are the amount of melanin concentrated in iris melanocytes, as well as the shape and distribution of melanosomes. Eye colour is highly variable in populations with European ancestry, in which eye colour categories cover a continuum of low to high quantities of melanin accumulated in the iris. A few polymorphisms in the *HERC2/OCA2* locus in chromosome 15 have the largest effect on eye colour in these populations, although there is evidence of other variants in the locus and across the genome also influencing eye colour. To improve our understanding of the genetic loci determining eye colour, we performed a meta-analysis of genome-wide association studies in a Canadian cohort of European ancestry (N= 5,641) and investigated putative causal variants. Our fine-mapping results indicate that there are several candidate causal signals in the *HERC2/OCA2* region, whereas other significant loci in the genome likely harbour a single causal signal *(TYR, TYRP1, IRF4, SLC24A4).* Furthermore, a short subset of the associated eye colour regions was colocalized with the gene expression or methylation profiles of cultured melanocytes *(HERC2, OCA2),* and transcriptome-wide association studies highlighted the expression of two genes associated with eye colour: *SLC24A4* and *OCA2.* Finally, genetic correlations of eye and hair colour from the same cohort suggest high pleiotropy at the genome level, but locus-level evidence hints at several differences in the genetic architecture of both traits. Overall, we provide a better picture of how polymorphisms modulate eye colour variation, particularly in the *HERC2/OCA2* locus, which may be a consequence of specific molecular processes in the iris melanocytes.

**Author Summary:** Eye colour differences among humans are the result of different amounts of melanin produced, as well as due to differences in the shape and distribution of the organelles in charge of producing melanin. Eye colour is a highly heritable trait, where several genes across the genome are involved in the process, but we currently do not fully understand which are the causal variants and how they modulate eye colour variation. By performing genome-wide association studies of eye colour across Canadian individuals of European ancestry, we identify several candidate causal signals in and near the gene *OCA2,* and one candidate signal in other genes, such as *TYR, TYRP1, IRF4* and *SLC24A4.* Furthermore, we provide insights about how significant loci may modulate eye colour variation by testing for shared signals with polymorphisms associated with the expression of genes and DNA methylation. Overall, we provide a better picture of the genetic architecture of eye colour and the molecular mechanisms contributing to its variation.

## Introduction

Pigmentation levels in the iris vary among humans, ultimately leading to different eye colours. The melanin pigment in the iris is synthesized in the melanocytes, within organelles named melanosomes (1). Eye colour diversity is a consequence of different amounts of melanin concentrated in the melanocytes of the iris. In addition, the shape and distribution of melanosomes influence eye colour variation. The mechanism is different from that of hair and skin pigmentation, in which two types of cells, melanocytes and keratinocytes (i.e. the epidermal melanin unit) play a key role in the production and distribution of melanin to give hair and skin colour (2–4). Additionally, out of the two types of melanin synthesized by melanocytes (i.e. eumelanin, a brown/black pigment and pheomelanin, an orange/yellow pigment), different categorical iris colours are a result of variation mainly on eumelanin content, whereas there is little, non-significant variation on pheomelanin quantity, based on measurements on cultured uveal melanocytes (5).

At a molecular level, blue irises appear as melanin-free melanocytes, in which molecules in the iris scatter short blue wavelengths to the surface (1). Green irises have medium levels of eumelanin, whereas high levels of eumelanin result in brown irises. Therefore, broad eye colour classifications (i.e. blue, green, hazel, brown) cover a continuum of low to high quantities of eumelanin accumulated in the iris (1). It is also well known that in some individuals there is a heterogeneous distribution of melanin in the iris, in which the peripupillary region is darker (e.g. richer in eumelanin) than the region closer to the sclera (1,6). This is known as iris heterochromia.

Twin studies have shown that eye colour is a highly heritable trait (> 85%) and that it does not significantly vary throughout an adult’s lifespan (7,8). Furthermore, association studies have demonstrated that eye colour has a polygenic architecture (9–12). Some of the loci with moderate/large effects associated with eye colour variation are at or near the following genes: *OCA2, TYR, TYRP1, SLC45A2, SLC24A4, SLC24A5* and *IRF4.* However, the variant with the largest effect on eye colour variation is an intronic SNP (rs12913832) located in an enhancer within the gene *HERC2* that regulates the expression of the downstream gene *OCA2* (13). Functional studies have shown that the T-allele of rs12913832 allows the formation of a chromatin loop with the promoter of *OCA2,* facilitating the transcription of the gene. In contrast, the C-allele hinders the formation of the chromatin loop, leading to a diminished expression of *OCA2* (13,14).

The SNP rs12913832 is the key regulatory element of *OCA2.* But it has been hypothesized that additional distal elements within the same region may be involved in the regulation of *OCA2,* a process that often is tissue-specific (14,15). In fact, through conditional analyses of association, genome-wide association studies (GWAS) have highlighted the presence of additional single-nucleotide polymorphisms (SNPs) associated with variation in pigmentary traits (i.e. skin, hair and eye pigmentation) within the *HERC2/OCA2* region (12,16–18).

These studies have identified variants within *HERC2* and *OCA2* that are in low (r^2^ < 0.2) linkage disequilibrium (LD) with rs12913832 (e.g. rs4778249, rs1667392, rs4778219, rs1800407, rs1448484), as well as other distant putative regulatory variants near or within the *APBA2* gene (e.g. rs4424881, rs36194177), which is located ~700kb away from *OCA2.* However, pinpointing additional causal variants within the *HERC2/OCA2* region is challenging due to the complex LD patterns among the genetic variants and the lack of tissue-specific regulatory annotations. For instance, the Gene and Tissue Expression (GTEx) database (19) includes skin tissue, which beyond a very small proportion of melanocytes, encompasses a diverse set of cell types not involved in pigmentation variation.

In order to improve our understanding of the genetic mechanisms behind eye pigmentation and melanocyte biology, in this paper we present the results of a GWAS of eye colour conducted in a Canadian cohort from the Canadian Partnership of Tomorrow’s Health (CanPath), along with fine-mapping analyses. We combined these results with gene expression and methylation data of cultured melanocytes by conducting colocalization analyses and transcriptome-wide association studies (TWAS). Our main results indicate that there are several candidate signals in the *HERC2/OCA2* region associated with eye colour, a different pattern from what is observed for hair colour in the same sampled population. By integrating expression and methylation data assayed in melanocytes, we gain a better picture about how genetic polymorphisms may modulate eye colour variation.

## Results

### Eye Colour Distribution in the CanPath Cohort

A total of 5,732 participants of the Canadian Partnership for Tomorrow’s Health (CanPath), who were genotyped using two genome-wide genotyping arrays (See Methods for details), also self-reported their natural eye colour using one of six possible answers: blue, grey, green, amber, hazel or brown. We excluded amber eye colour individuals due to the low number of individuals who self-reported this category. After quality control of the genotypes (i.e. exclusion of poor-quality samples and PCA outliers), we kept 5,641 individuals for further analyses. The distribution of eye colour categories was similar across all provinces sampled (Figure 1), with green and hazel being the least frequent categories and blue being the most common one. One exception is the significantly low proportion of individuals who self-reported blue eye colour in Quebec, compared to other provinces (chi-square test: 104.39, df = 4, p-value < 0.01). This pattern may be explained by the high proportion of French ancestry in the Quebec population (Supplementary Figure 1) due to the migration and settlement of French people in the province relatively recently (20). Additionally, a higher proportion of females self-reported green and hazel eye colours, relative to their male counterpart (chi-square test for green and hazel combined = 244.49; df = 1; p-value < 0.01), which is similar to the observations previously reported in the case of green eye colour (9).

**Figure 1.**
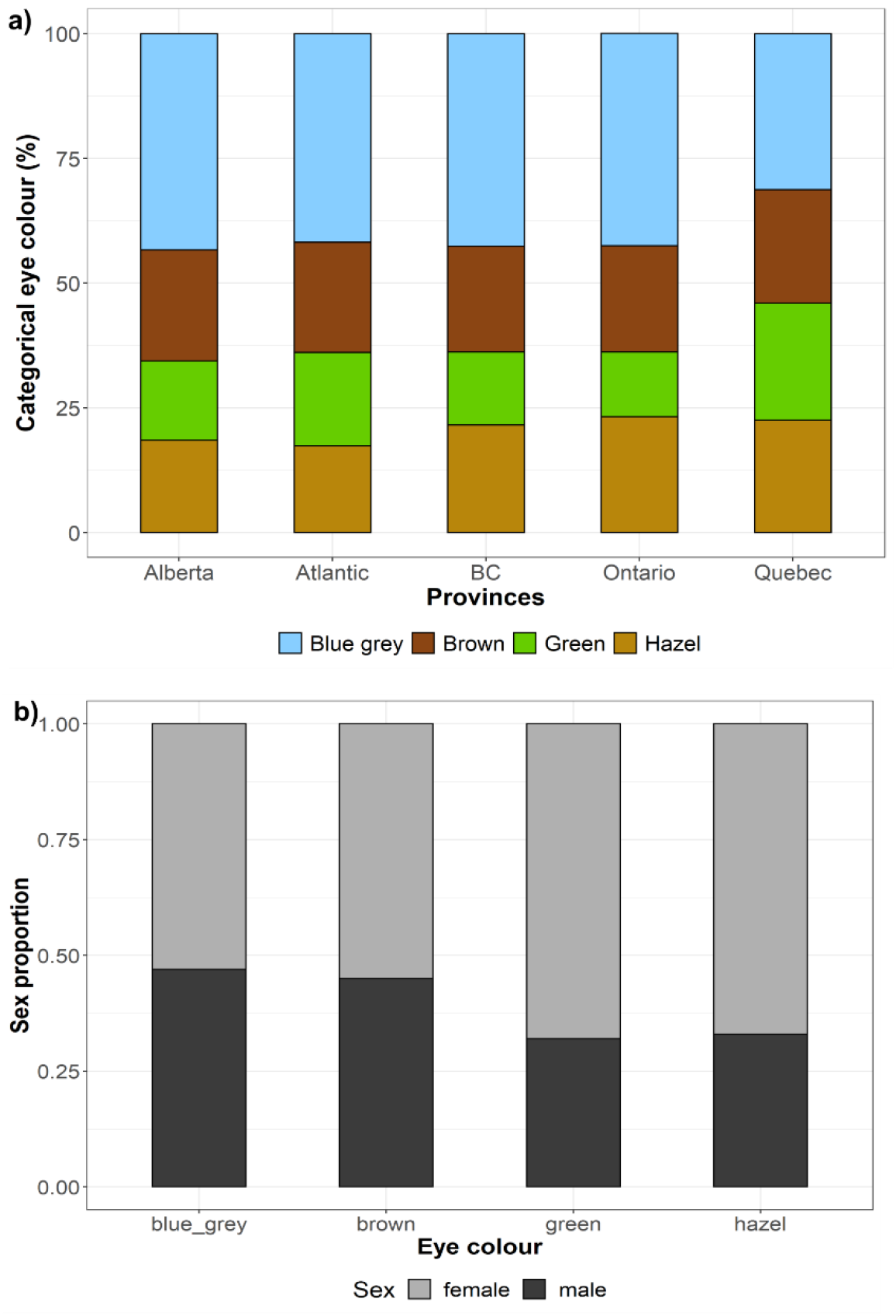
Distribution of eye colour categories in the CanPath a) Percentage of the hair colour categories stratified by province, **b)** Proportion of sexes across eye colour categories.

### Genome-wide Association Studies and Meta-Analyses

We performed GWAS of eye colour on each genotyping array (genotyped and imputed single-nucleotide polymorphisms (SNPs)) using a linear mixed model and an additive genetic model, using GCTA 1.26.0 (21,22). We coded eye colour categories as follows: 1 = blue or grey, 2 = green, 3 = hazel and 4 = brown.

We included sex, age and the first ten principal components (PCs) as fixed effects and a genetic relationship matrix (GRM) as random effect to control for subtle population structure. We did not detect residual population substructure, based on Q-Q plots, in which observed p-values did not show an early deviation from the expected p-values (Supplementary Figure 2).

We then carried out a meta-analysis using the summary statistics (log of odds ratio and standard error) including the two GWAS on METASOFT v2.0.1 (23). Q-Q plots of the meta-analyses (Supplementary Figure 3) and LD Score regression (intercept = 0.9935), indicated no residual population structure. We identified several known genome-wide significant loci (p-value ≤ 5e-08) associated with eye colour (Figure 2; Supplementary Figure 4), overlapping or near the genes *TYRP1* (lead SNP: rs1326779; beta = 0.139; SE = 0.024)*, IRF4* (lead SNP: rs12203592; beta = −0.164; SE = 0.029), *TYR* (lead SNP: rs1126809; beta = −0.136; SE = 0.024), *SLC24A4* (lead SNP: rs4144266; beta = −0.124; SE = 0.022) and *HERC2* (lead SNP: rs1129038; beta = −1.239; SE = 0.024). Additionally, we observed a signal on chromosome 6 overlapping the *ILRUN* gene (lead SNP: rs116072038; beta = −0.438; SE = 0.077), a locus that has not been previously associated with pigmentation. Supplementary File 1 summarises the suggestive and genome-wide associated SNPs.

**Figure 2.**
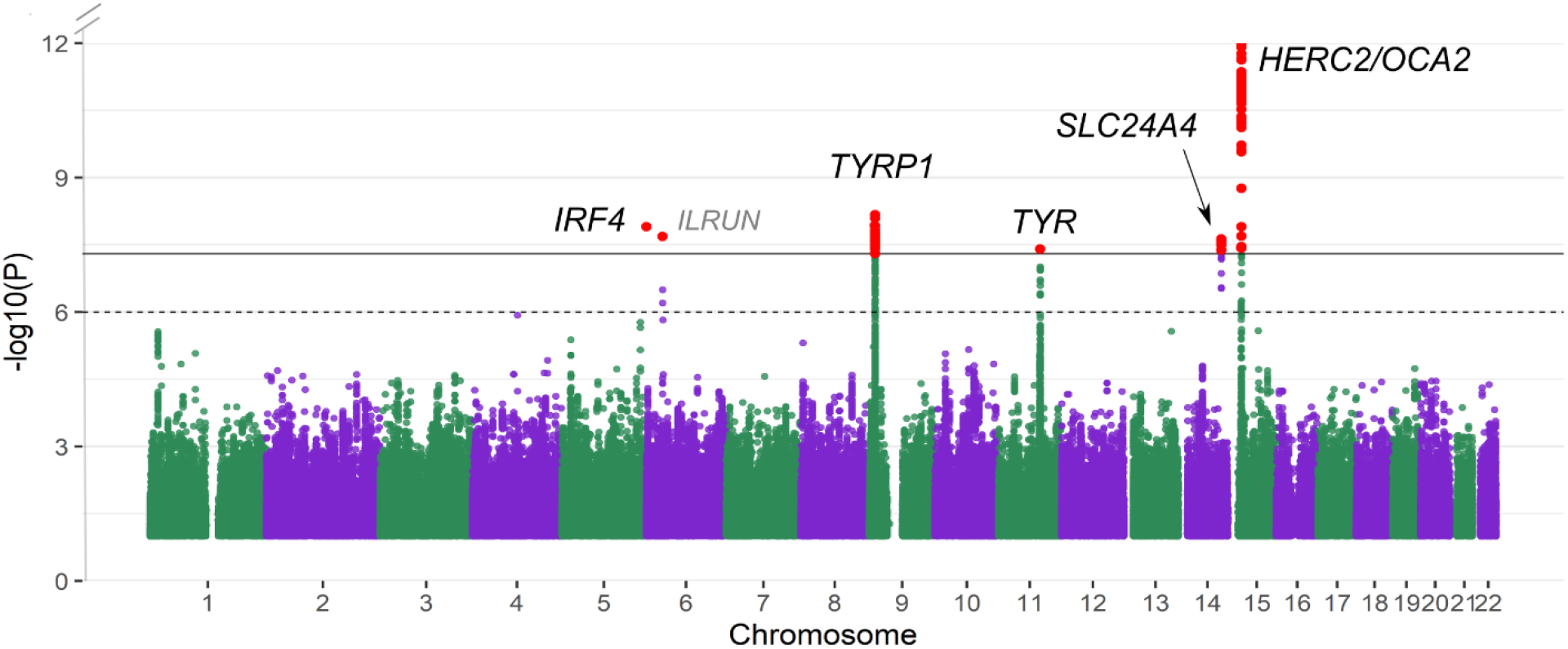
Manhattan plot of eye colour meta-analysis based on a linear mixed model. The dotted line indicates the suggestive threshold (p = 1e-06) and the continuous line denotes the genomewide threshold (p = 5e-08). The Y-axis has been limited to truncate strong signals at the locus in chromosome 15. The full figure is available as Supplementary Figure 3.

### Fine-mapping of GWAS hits

We conducted approximate conditional and joint analyses of association using GCTA-COJO (24), to investigate if the genome-wide significant loci were being driven by one or more independent signals.

On the *IRF4, TYRP1, TYR* and *SLC24A4* regions, we identified one independent genome-wide significant SNP per locus (Supplementary Table 2), corresponding to known causal variants, such as rs12203592 on *IRF4* and rs1126809 on *TYR*. The lead SNP on *SLC24A4* is in high LD (r^2^ = 0.98) with rs12896399, a SNP previously associated with pigmentation (9,25). In the case of *TYRP1,* the selected SNP (rs1326779) is downstream of *TYRP1* and has not been previously highlighted in eye colour studies. On the *HERC2/OCA2* region, we identified six independent SNPs overlapping *OCA2* and *HERC2* with p-values in the conditional analysis exceeding the genome-wide significant threshold (Supplementary Table 2). Several of these SNPs have evidence of heterogeneity among the two studies, as indicated by I^2^ and Cochran’s Q values in the meta-analysis. However, they all have genome-wide significant p-values in the random effects (RE2) model too, which takes into account heterogeneity among studies (Supplementary Table 2). To validate our results, we carried out the same GCTA-COJO analysis a second time, using the GSA array as LD reference. We obtained concordant results, with multiple independent SNPs on the *HERC2/OCA2* region and a single SNP highlighted on the other pigmentation associated loci (i.e. *IRF4, TYR, SLC24A4* and *TYRP1*) (Supplementary Table 3).

We also carried out a Bayesian fine-mapping analysis, in which all possible combinations of SNPs are iteratively considered without arbitrary selection of conditioned SNPs. We used the program FINEMAP (26) to perform fine-mapping analysis and to identify candidate causal SNPs for functional prioritization. In agreement with the GCTA-COJO analysis, by using FINEMAP we identified known pigmentation genes harbouring one causal signal within *IRF4, TYR, SLC24A4* and *TYR* (Supplementary File 2). On the *IRF4* locus, the only candidate causal SNP with considerable evidence of causality (log_10_BF > 2) was the same SNP highlighted by GCTA-COJO (rs12203592; PIP = 0.999). In contrast, the missense SNP rs1126809 on *TYR* had a low posterior inclusion probability (PIP = 0.209), due to high LD with other nearby SNPs. Other candidate causal SNPs in the 95% credible set of the *TYR* locus include intergenic variants and one SNP (rs11018578) on the 3’UTR region of *NOX4.* On the *SLC24A4* locus, the variants with considerable evidence of causality (i.e. log_10_BF ≥ 2) include intronic SNPs within *SLC24A4* and other variants upstream of the gene, including the SNP rs12896399 (log_10_BF = 2.58) (Supplementary File 2).

On the *TYRP1* locus, all candidate causal SNPs in the credible set had a low (< 0.1) PIP, most likely due to high LD among multiple SNPs in the locus (Supplementary Figure 5). The 95% credible set includes rs10809826 and rs1408799, two SNPs which have been previously associated with eye colour (12,27–29). Amongst the SNPs with log_10_BF > 2 in the same 95% credible set, rs13297008 is located upstream of *TYRP1* and it overlaps a DNase Hypersensitive Site identified in foreskin melanocytes (Supplementary File 2), indicative of an active transcription region. Given that some genetic variants may not be present in our dataset due to poor or lack of imputation, we further explored the regulatory annotation of SNPs within the same LD block as rs13297008 (r^2^ ≥ 0.8) using HaploReg (version 4) (30,31). We observed that rs13297008, rs2733831 and rs13296454 are in an active transcription start site (TSS) state on foreskin melanocytes only (across the tissues tested with the 15-core chromatin states). These SNPs are either suggestive of association or genome-wide significant, and all three are within the 95% credible set (Supplementary File 1).

### Multiple HERC2/OCA2 variants associated with eye colour variation

By applying a Bayesian fine-mapping approach on the *HERC2/OCA2* region, we identified five putative causal signals (i.e. five 95% credible sets) associated with eye colour. Within these signals, three SNPs had a PIP > 0.98 (Table 1; Figure 3A). These results suggest independent causality of the various signals in the locus. One of the candidate SNPs within *HERC2* is rs12913832, a known enhancer that regulates the expression of *OCA2* (13). In addition, three other SNPs within *HERC2* and one within *OCA2* were nominated as candidate causal loci, all of which fall within introns. These results are similar to the conditional analysis with GCTA-COJO, in which the independent SNPs in the locus encompass both *OCA2* and *HERC2,* but the selected SNPs do not fully overlap. Importantly, all fine-mapped SNPs in the locus had genome-wide significant p-values on each sample and the same direction of effect. We annotated the putative regulatory function of the SNPs in all five credible sets using diverse databases (e.g. ENCODE, Roadmap Epigenomics Project). Aside from the overlap of rs12913832 *(HERC2)* with an open chromatin region in foreskin melanocytes, only the SNP rs117007668 is located within an open chromatin region in foreskin melanocytes (Supplementary File 2).

**Figure 3.**
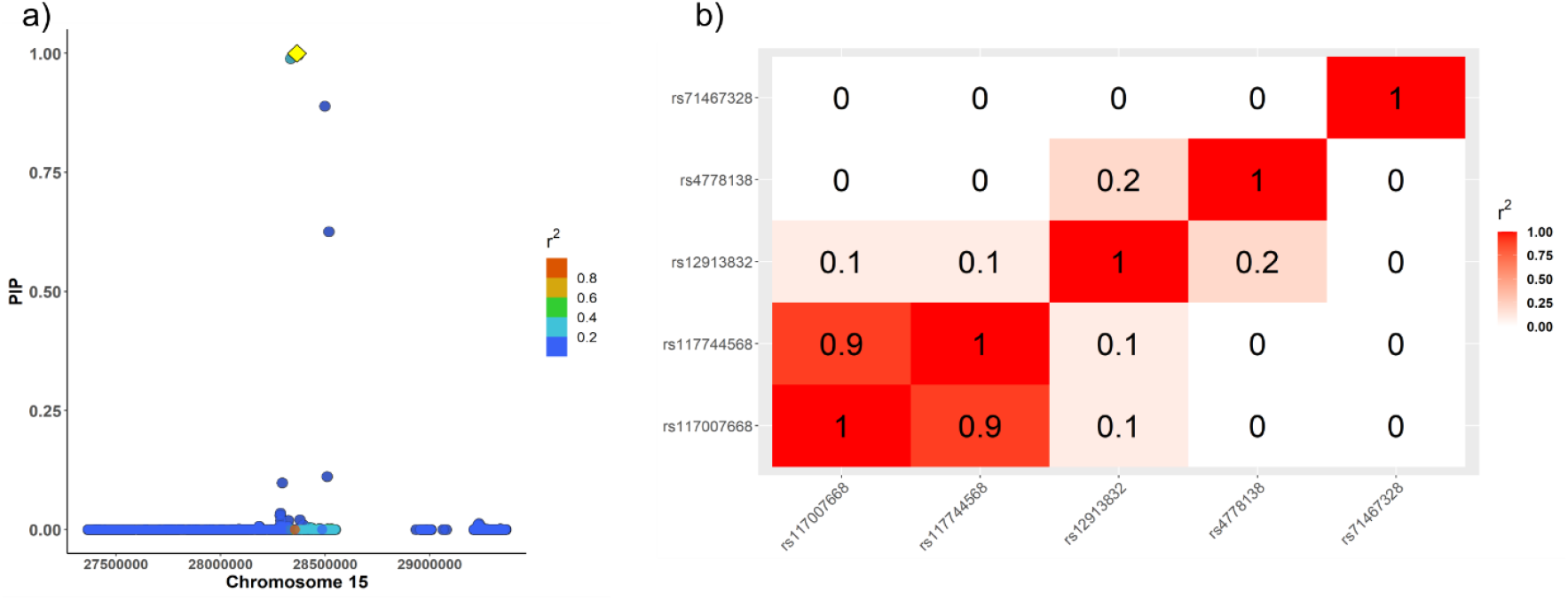
Fine-mapping of the *HERC2/OCA2* locus (chromosome 15) associated with eye colour, a) FINEMAP regional plot of the posterior inclusion probability (PIP), in which the lead SNP (rs12913832) is highlighted in yellow and LD (r^2^) correlations are shown in respect to the lead SNP. b) Matrix of LD correlations among the SNPs with the highest PIP on each of the five 95% credible set.

**Table 1.** Summary statistics of the candidate causal SNPs with log_10_BF ≥ 2 in the *OCA2/HERC2* region on chromosome 15, associated with eye colour. **FE = fixed-effects model; RE2= random-effects model; MAF = minor allele frequency; I^2^ and Cochran’s Q: meta-analysis heterogeneity indices; PIP = posterior inclusion probability; log_10_BF = log_10_ of Bayes Factor.**

| Credible Set | rsid | Position in chr 15 | Gene | Beta | SE | p-value (FE) | p-value (RE2) | MAF | $I^2$ | Cochran's Q | Cochran's Q p-value | PIP | $\log_{10}\text{BF}$ |
| --- | --- | --- | --- | --- | --- | --- | --- | --- | --- | --- | --- | --- | --- |
| 1 | rs12913832 | 28365618 | *HERC2* | -1.26 | 0.02 | 0 | 0 | 0.23 | 96.25 | 26.66 | 2.43E-07 | 1.00 | 13.14 |
| 2 | rs117007668 | 28371422 | *HERC2* | 0.99 | 0.08 | 5.43E-38 | 8.30E-38 | 0.01 | 81.03 | 5.27 | 0.02 | 1.00 | 5.59 |
| 3 | rs4778138 | 28335820 | *OCA2* | 0.81 | 0.03 | 2.56E-162 | 1.51E-161 | 0.13 | 0.00 | 0.25 | 0.62 | 0.99 | 5.08 |
| 4 | rs117744568 | 28498692 | *HERC2* | 0.87 | 0.06 | 3.95E-46 | 1.01E-45 | 0.02 | 73.99 | 3.85 | 0.05 | 0.89 | 4.04 |
| 4 | rs117743506 | 28510460 | *HERC2* | 0.85 | 0.06 | 6.46E-46 | 1.96E-45 | 0.02 | 65.92 | 2.93 | 0.09 | 0.11 | 2.23 |
| 5 | rs71467328 | 28518229 | *HERC2* | -0.48 | 0.05 | 5.60E-18 | 1.13E-17 | 0.04 | 27.37 | 1.38 | 0.24 | 0.63 | 3.36 |
| 5 | rs1597196 | 28294922 | *OCA2* | 0.42 | 0.03 | 3.86E-55 | 1.27E-54 | 0.18 | 63.25 | 2.72 | 0.10 | 0.10 | 2.17 |

We then explored the LD patterns among the candidate causal variants (Table 1) in the credible sets using the CanPath genotypes (i.e. the same LD matrix used for fine-mapping) and considering the genotype probabilities, computed with LDStore v2.0 (32). Amongst the top candidate causal variants across the five credible sets (Table 1), most correlations are low (r^2^ ≤ 0.2), with the exception of rs117007668 and rs117744568, which are in high LD (r^2^ = 0.9) (Figure 3B). We further compared D’ values among these same SNPs computed on LDLink, using as a proxy the European populations of the 1000 Genomes Project (33). The D’ patterns, compared to r^2^ values, reflect the allele frequency differences among the SNPs (D’ > 0.6 in most cases) and suggest that the candidate causal SNPs are not in complete linkage equilibrium (Supplementary Figure 6).

Finally, we explored with HaploReg (version 4) (30,31) if other variants in the same LD block as our credible set SNPs in the *OCA2/HERC2* locus harbour a putative regulatory function, to consider genetic variants that may have not been present in out dataset after imputation. By using as input the top SNP on each of the five credible sets (Table 1), we identified a nominally significant enrichment of enhancers (as defined by the 15-state core ChromHMM model) in foreskin melanocytes (binomial test compared to all 1KGP variants with MAF ≥ 5%; p-value = 0.0169). The SNP rs117743506, which is in high LD with rs117744568 and in the same credible set, has an enhancer state in foreskin melanocytes. Additionally, it may alter the motif of POU3F2, a transcription factor (TF) present in melanoma-cell lines known to alter the expression of pigmentation genes (i.e. *MITF, KITLG),* although a recent study suggests that this TF does not have a role in normal skin melanocytes (34,35).

### Associations with eye colour in recent studies

By investigating eye colour variation in the *OCA2* locus, Andersen *et al.* (2016) identified that two missense SNPs within *OCA2* (rs121918166 and rs74653330) have a measurable effect on eye colour variation, in which the alternative alleles decrease melanin levels, even in a heterozygote state. These two SNPs were rare in our sample (MAF < 1%), therefore we did not consider them in our GWAS analyses. However, we identified in our sample a subset of individuals (N= 904) harbouring the rs12913832:GG genotype, 21 of which self-reported brown eye colour, 631 green eye colour and 252 hazel eye colour, suggesting that the rs12913832:GG genotype does not exclusively yield a blue eye colour.

Adhikari *et al.* (2019) conducted conditional analyses of association of eye colour (measured qualitatively and quantitatively) in Latin American individuals with mainly European and Native American ancestry and identified up to five independent signals in the *HERC2/OCA2* locus (indexed by: rs4778219, rs1800407, rs1800404, rs12913832 and rs4778249). Aside from rs12913832, none of their index SNPs are within the candidate causal SNPs in our sample, even though rs1800404 was genome-wide significant in our metaanalysis (p-value = 1.18e-11). Additionally, they identified three novel loci associated with eye pigmentation: *DSTYK* (chromosome 1)*, WFDC5* (chromosome 20) and *MPST* (chromosome 22). We followed up each of the three index SNPs in our meta-analyses (rs3795556, rs17422688 and rs5756492), but failed to replicate the former two SNPs using a Bonferroni correction (p-value threshold = 0.005), whereas rs5756492 was not present in our meta-analysis. These differences may be driven by population ancestry differences, given that the CANDELA cohort includes recently admixed individuals from Latin America, although the phenotyping approach may also be driving these differences.

Finally, the largest eye colour GWAS to date conducted in populations of mainly European ancestry reported several novel loci, which had not been previously associated with eye colour, and a subset of them has not been previously associated with any pigmentation traits (eye, hair or skin pigmentation) (37). However, the *ILRUN* locus identified in the present study was not amongst their novel signals, nor we were able to replicate it. Additionally, they conducted conditional analyses of association to identify secondary signals on the significant loci, in which they identified a total of 115 independent signals, including three signals in chromosome X. Notably, they identified 33 independent signals in the *HERC2/OCA2* region and two signals in the nearby gene *GABRB3.* We followed-up their signals (Supplementary Table 1 from (37)) in our meta-analyses, and identified 50 SNPs nominally significant in our meta-analysis (p-value ≤ 0.05), all with a consistent direction of effect between both studies (Supplementary Table 4). This set of SNPs includes novel associations with eye colour and/or pigmentation traits included in their study (i.e. *DTL, MITF, PDCD6/AHRR, ADRB2, GCNT2* and *SIK1).* After considering a Bonferroni correction (0.05/112; p-value ≤ 4.46e-04), 16 SNPs remained significant, all of which overlap known pigmentation genes (*IRF4, TYRP1, TYR, OCA2* and *HERC2*).

Lastly, we checked if the independent SNPs associated with eye colour identified with GCTA-COJO by Simcoe *et al.* (37) overlap with our eye colour SNPs highlighted by GCTA-COJO (Supplementary Table 2). We identified an overlap of three SNPs: rs12203592 (*IRF4*), rs1126809 (*TYR*) and rs1129038 (*HERC2*). Interestingly, even though Simcoe *et al.* identified several independent loci in the *OCA2/HERC2* region, the known rs12913832 SNP is not among them, likely because another SNP in perfect LD has a lower p-value, which is similar to what we observe in our GCTA-COJO analysis (i.e. rs1129038). We also compared the same independent SNPs identified by Simcoe *et al.* with our candidate causal loci as defined by FINEMAP, and found three overlapped SNPs: rs12203592 *(IRF4),* rs1126809 (*TYR*) and rs13297008 (*TYRP1*).

### Colocalization with Expression and Methylation QTLs from Cultured Melanocytes

We performed colocalization analyses with hyprcoloc (38) of the eye colour meta-analysis with melanocyte gene expression and methylation *c/s*-QTLs (eQTLs, meQTLs, respectively) to explore if there were shared causal signals (See Methods for details). Through the colocalization of GWAS with eQTLs, we identified a region overlapping *OCA2* and AC090696.2 (the latter being a transcript which partially overlaps *OCA2)* (Table 2), in which the candidate marker is rs12913832. We also colocalized meQTLs with GWAS hits (Table 2) overlapping the gene body of *HERC2* (tagged by cg25622125, cg27374167 and cg05271345) using a posterior probability threshold of 0.8. Notably, we did not find colocalized eQTLs on the *TYRP1* locus that passed the probability cuttof.

**Table 2.** Colocalization results of expression and methylation QTLs (eQTL and meQTL, respectively) with GWAS eye colour SNPs, showing colocalized SNPs with a posterior probability ≥ 0.8. The methylation annotation indicates the location with respect to the nearest gene (TSS = transcription start site), as well as the location of the tagged CpG marker within the CpG island.

| Chromosome | Candidate SNP | Posterior Probability | Regional Probability | Posterior explained by SNP | Gene/Methylation Annotation | QTL |
| --- | --- | --- | --- | --- | --- | --- |
| 15 | rs12913832 | 0.9999 | 1 | 1 | *OCA2* | eQTL |
| 15 | rs12913832 | 0.9864 | 0.9891 | 1 | AC090696.2 | eQTL |
| 15 | rs12913832 | 0.9991 | 0.9995 | 1 | *HERC2* (Body) \| OpenSea | meQTL |
| 15 | rs12913832 | 0.9855 | 0.9872 | 1 | *HERC2* (Body) \| S_Shelf | meQTL |
| 15 | rs12913832 | 0.9807 | 0.9831 | 1 | *HERC2* (Body) \| S_Shore | meQTL |

### Transcriptome-Wide Association Studies (TWAS)

We conducted TWAS using a subset of the CanPath cohort as LD reference, and the expression weights from cultured melanocytes to predict the gene expression profile with FUSION (39). Our results highlighted the expression of three genes as significantly associated with eye colour: *OCA2, SLC24A4* and *RIN3* (Supplementary Table 5; Supplementary Figure 7). The gene *RIN3* is located near *SLC24A4,* and by conducting conditional TWAS we have shown that these two genes are not independent from each other (Supplementary Figure 8).

### Genetic Correlations

We calculated the genetic correlation between eye and hair colour using the data from the two CanPath genotyping arrays for which we had full phenotype data. Using a linear scale for both traits and the same covariates as used in the GWAS (See Materials and Methods for details), there is a genetic correlation (rg) of 55% (SE = 0.12; p-value = 7.33e-6) and 69% (SE = 0.21; p-value = 0.001) on the UK Biobank and GSA arrays, respectively. Similar to the approach used in a previous study (40), we then calculated the genetic correlation without controlling for the effect of significant principal components, and obtained genetic correlation values of 63% (SE = 0.08; p-value = 3.41e-15) and 79% (SE = 0.15; p-value = 1.39e-07) on the UK Biobank and GSA arrays, respectively. These results are in line with the genetic correlations previously reported (40), in which they found a lower correlation when including principal components as covariates, due to the correlations between ancestry captured by the PCs and eye or hair colour. Nevertheless, by not controlling for significant PCs, we may as well be capturing ancestry in the genetic correlation estimation.

### Eye Colour as a Risk Factor for Uveal Melanoma

Uveal melanomas (UM) are rare cancers that arise from melanocytes in the pigmented uveal tissues of the eye (i.e. iris, ciliary body and choroid) (41). One of the risk factors for UM is iris colour, with light eye colours (e.g. blue) associated more frequently with UM. It has also been described that iris melanomas occur more frequently in the inferior quadrant of the iris compared to other parts of the eye. This is hypothesized to be due to the higher exposure to UV in that quadrant, followed by the high phototoxic effect of pheomelanin in the case of low eumelanin content (5,41–43). However, a study suggested that the DNA damage signature of most UM cases does not correspond to that of UV damage, suggesting that UV radiation is not a significant factor for UM (44). Additionally, it has also been shown that in the absence of high eumelanin levels, pheomelanin generates more reactive oxygen species independent of UV light (45).

Only a few studies have investigated the genetic basis of uveal melanoma, identifying risk variants in pigmentation SNPs in *HERC2, OCA2* and *IRF4* (46), the loci with the largest effects on eye colour in the present study. Based on the GWAS Catalog search term ‘uveal melanoma’, only two studies have been reported, which have been conducted with moderate sample sizes (47,48). We followed-up 35 unique SNPs associated with UM (p ≤ 1e-6) in our meta-analysis (Supplementary Table 6) and found that none of them are significantly associated with eye colour, using a Bonferroni corrected p-value = 0.001. Of note, the *HERC2* rs12913832 SNP is not genome-wide significant in neither UM GWAS and the only SNP suggestive of association in the *HERC2/OCA2* region is rs11074306. Future UM studies with larger sample sizes may increase the power to associate other pigmentation loci with uveal melanoma risk that have small effects, similar to what is observed for cutaneous melanoma. It would be relevant to characterize the differences among the UM subtypes, to be able to evaluate the risk of UV radiation and eye colour variation in UM susceptibility.

## Discussion

In this paper we present the results of our genome-wide association studies of eye colour, as measured categorically through self-reports, from 5,641 participants of the Canadian Partnership for Tomorrow’s Health (CanPath). We did not identify new loci associated with eye colour that were successfully replicated, and we focused on performing downstream analysis to pinpoint candidate causal SNPs, specifically on those loci for which a functional variant has not yet been identified, or in which there is evidence of more than one independent signal. We found that fine-mapping provides evidence for multiple independent SNPs within the *HERC2/OCA2* region, whereas other loci likely have a single causal signal. Furthermore, we characterized our GWAS signals by using colocalization analyses with expression and methylation QTLs of cultured melanocytes, and conducted TWAS, in which we identified the expression of *SLC24A4/RIN3* and *OCA2* as significantly associated with eye colour. Lastly, we explored the genetic correlations between hair and eye colour in the CanPath cohort.

One of the caveats of this study is that we utilized eye colour categories self-reported by participants of the CanPath cohorts as categorical classifications do not capture as well iris colour variation as quantitative measures (11,49,50). In our sample we have identified significant differences in self-reporting of eye colour between sexes, and although these may be a reflection of true sex differences, as has been previously reported in other studies reporting self-assessed categorical eye colour (9), we cannot discard the possibility of a self-reporting bias. This limitation is counter-balanced by the relatively large sample size, in comparison to the majority of previous studies (16,51,52), with the exception of the largest recent GWAS of eye colour (37). Furthermore, significantly associated loci from self-reported eye colour are useful in forensics, for predicting eye colour categories, which the human eye can easily distinguish. Indeed, we have here identified most signals associated with eye colour that are used in the IrisPlex eye colour prediction system (53–55). However, it is important to point out that structural features of the iris (i.e. contraction furrows, Wolfflin nodules, heterochromia) also contribute to colour perceptions, but we are not able to distinguish them using the current dataset.

Through our GWAS meta-analysis we identified five known loci associated with eye colour, encompassing the genes *SLC24A4, IRF4, TYRP1, TYR* and *HERC2/OCA2,* similar to what was identified in a recent large GWAS of both categorical and quantitative eye colour loci in a Latin American (CANDELA) cohort (12). The only two known pigmentation loci that our GWAS failed to identify as significantly associated with eye colour encompass the genes *SLC24A5* on chromosome 15 and *SLC45A2* on chromosome 5, in which missense variants (rs1426654 and rs16891982) are known to alter pigmentation traits (56). The missense SNP on *SLC24A5* was rare in our sample (MAF < 1%) hence excluded, in line with frequencies observed in the 1000 Genomes Project European populations, in which the alternative allele is nearly fixed. In the case of the *SLC45A2* locus, the missense SNP did not reach genome-wide significance (p-value = 1.71e-6).

Our fine-mapping analyses identified known causal pigmentation loci in the credible sets, such as rs12203592 on *IRF4,* rs1126809 on *TYR* and rs12913832 on *HERC2.* Contrary to what has been observed for hair pigmentation (12), we identified here one independent SNP in the *TYR* locus associated with eye colour, even though there is at least another independent missense variant (rs1042602) known to alter melanin synthesis within the same gene, in addition to putative regulatory variants in the upstream *GRM5* gene associated with skin pigmentation (9,12,16,17,50,57–59). This result is also in line with what was reported in the CANDELA study (12), in which, compared to hair colour, they identified a single candidate SNP in the *TYR* locus associated with eye colour. This exemplifies the importance of characterizing the genetic architecture of different pigmentation traits independently and opens up new questions to investigate the different mechanisms involved in melanin synthesis between cutaneous vs. iris melanocytes.

We conducted TWAS and colocalization analysis with expression and methylation QTLs to further explore the shared causal signals among these phenotypes. Through colocalization with melanocyte eQTLs, we found colocalization in the *OCA2* region, regulated likely by the SNP rs12913832 in the nearby *HERC2* gene. Similarly, colocalization with meQTLs highlighted a signal in the *HERC2* locus. The shared signals between meQTLs, eQTLS and eye colour GWAS hits in the *HERC2* region may suggest that DNA methylation could play a role in the differential expression of *OCA2,* thus influencing the eye colour phenotype, although this cannot be confirmed with the current evidence. Further analyses, such as Mendelian randomization, will be useful to evaluate causal associations among these traits (e.g. Bonilla *et al.,* 2020).

We did not find colocalization of GWAS SNPs with eQTLs on the *TYRP1* locus, even though our GWAS and fine-mapping results suggest a regulatory role of the candidate causal variants in this locus due to: 1) the location ~11kb upstream of the gene, and 2) the overlap of a SNP (rs13297008) with open chromatin regions in foreskin melanocytes. Additionally, this gene was absent from the TWAS expression weights dataset, suggesting that the gene expression in the current dataset is not sufficiently heritable (i.e. heritability p > 0.01). Therefore, we are not able to provide evidence of the mechanism in which the variants in the locus affect pigmentation variation, nor we are able to nominate a single causal SNP.

In the *TYRP1* locus there is one likely candidate of causality (i.e. log_10_BF > 2 combined with regulatory annotation): rs13297008, which is in the same credible set as two previously associated SNPs with hair and eye colour in the same locus (rs1408799 and rs10809826) (12,25). However, neither rs1408799 nor rs10809826 had considerable evidence of causality (log_10_BF < 2) in our analysis. In contrast, by further exploring regulatory annotations of markers in LD, we prioritized two other putative functional SNPs near the TSS of *TYRP1* (rs2733831 and rs13296454). Of note, HaploReg (version 4) uses the 1000 Genomes Project Phase I data, therefore there may be missing variants, or variants present but not accurately assessed. Nonetheless, this finding shows the importance of fine-mapping GWAS loci along with diverse annotations, which could provide further evidence to prioritize candidate SNPs based on their putative function in a tissue-specific manner, in this case a possible regulatory function in melanocytes. Additionally, it is important to note that the IrisPlex eye colour prediction algorithm does not include SNPs within the *TYRP1* locus, even though it is one of the highly associated loci in our results.

The cultured melanocyte expression and methylation QTLs we used for colocalization and TWAS come from newborn foreskin melanocytes (61,62). Similarly, the regulatory annotations from the ENCODE and Roadmap Epigenomics projects (63,64) also come from melanocytes, keratinocytes and fibroblasts from skin tissue. The melanocytes from skin and iris have several similarities and same embryological origin, but there are also significant differences between them. For instance, the melanosomes within the iris melanocytes are retained in the cytoplasm and they are not transferred through dendrite-like structures to adjacent keratinocytes, as it is the case in the skin and hair melanocytes (1). Moreover, the iris melanocytes are not reactive to the alpha melanocyte stimulating hormone (α-MSH) (65) and instead, alternative signaling cascades trigger and regulate melanogenesis (66). Therefore, future QTL efforts using a more precise tissue type, such as uveal melanocytes, may aid in characterizing the regulatory differences between cutaneous and iris melanocytes.

The *HERC2/OCA2* region on chromosome 15 has the strongest effect on eye colour variation, such that blue eye colour was initially considered a Mendelian trait (1,67). Mutations on *OCA2* are known to cause oculocutaneous albinism II, but less deleterious mutations result in a decrease of eumelanin by increasing the acidity of the melanosomes and leading to a suboptimal performance of tyrosinase (68–71). The most significant variant associated with blue vs. brown eye colour is rs12913832, an enhancer of the expression of *OCA2* (13), while the same polymorphism only causes a mild decrease of hair and skin eumelanin content, suggesting that the effect of this locus is different between dermal and iris melanocytes (14).

Our findings suggest that it is likely that other SNPs in the locus also have an effect on the expression of *OCA2* in the iris. For instance, a subset of participants harboured the rs12913832 homozygous genotype associated with blue eye colour (i.e. GG), but they self-reported non-blue eye colour. Therefore, the expression of *OCA2* might be induced by other regulatory variants in the locus, counteracting the effect of rs12913832, as has been previously proposed (36). An alternative explanation could be that a subset of participants self-reported their eye colour inaccurately, a hypothesis that we are not able to discard. Nonetheless, through our fine-mapping analyses we have nominated additional putative regulatory variants that may also be modulating the expression of *OCA2.*

Additionally, it is possible that genetic interactions between *IRF4* and *OCA2* also play a role. For instance, it is known that individuals may have blue eye colour when harbouring one or two rs12913832 A-alleles *(HERC2),* associated with non-blue eye colour, along with one or two rs12203592 T-alleles, associated with light eye colour (72). Similarly, it has been recently suggested that SNPs in the genes *TYR* (rs1126809)*, TYRP1* (rs35866166, rs62538956) and *SLC24A4* (rs1289469) may be responsible for the brown eye colour in individuals of European ancestry with a rs12913832 homozygous G-allele background (73).

Finally, genetic correlations among hair and eye colour in the CanPath cohort are high, in line with what has been previously reported (40), and considering the effect that most genes have in both phenotypes too (e.g. *SLC24A4, IRF4, OCA2).* However, we have demonstrated that certain genetic differences come to light when investigating candidate causal variants across the genome. Within the CanPath cohort, we observed that red hair colour is driven mainly by multiple candidate causal signals in the *MC1R* locus, and that variants within the same gene also have a significant effect upon blonde hair colour. In contrast, variants within *MC1R* and its antagonist, *ASIP,* are not associated with eye colour, which may be explained by the fact that *MC1R* is not expressed in iris melanocytes (65). Additionally, this may explain why iris melanocytes do not respond to UV radiation as opposed to skin melanocytes. *HERC2/OCA2* is the most significant locus in our analysis of blond vs. black and brown vs. black hair color (although as described above, *MC1R* is the most important locus determining red hair colour). *HERC2/OCA2* is also the most significant locus for eye colour. However, the signal from hair colour is primarily driven by rs12913832, whereas there are several independent signals in *HERC2/OCA2* associated with eye colour. Lastly, even though *IRF4,* a transcription factor that upregulates tyrosinase, has a large effect on both blonde hair and blue eye colour, the direction of effect of the causal SNP rs12203592 is opposite for both traits: the derived T-allele is associated with blue eye colour, whereas the same allele is associated with the presence of brown hair colour (74).

## Materials and Methods

### Canadian Partnership for Tomorrow’s Health Participants

This study was approved by the University of Toronto Ethics Committee (Human Research Protocol # 36429) and data access was granted by the Canadian Partnership for Tomorrow’s Health (Application number DAO-034431). The samples in this study correspond to a subset of 5,675 individuals from the Canadian Partnership for Tomorrow’s Health (CanPath), which were sampled in different provinces: Alberta (N= 926; 16.4%), Atlantic Coast Provinces (i.e. New Brunswick, Newfoundland, Nova Scotia and Prince Edward Island) (N= 385; 6.8%), British Columbia (N= 965; 17.1%), Ontario (N= 934; 16.5%) and Quebec (N= 2434; 43.1%). We selected the individuals who self-reported having European-related ancestry and for whom self-reported eye colour was available (N= 5,641).

### Genotyping of Participants and Quality Control

Individuals who self-reported as having European-related ancestry were genotyped between 2012 and 2018 using two different genotyping array chips: Axiom 2.0 UK Biobank (Affymetrix) (N= 3,212) and the Global Screening Array (GSA) 24v1+MDP (N= 2,429) by the Canadian Partnership for Tomorrow’s Health (CanPath). The number of single nucleotide polymorphisms (SNPs) of these chip arrays ranges between 658,296 and 813,168 SNPs.

We performed genotype quality control for each array chip separately by first filtering out variants that deviated in minor allele frequency > 0.2 from the 1000 Genomes Project Phase 3 European sample (1KGP-EUR), GC/TA variants with minor allele frequency > 0.4 in the 1KGP-EUR and flipping alleles according to the 1KGP-EUR, using a Perl script (version 4.2) (75). Afterwards, we used PLINK (version 1.9) (76,77) to filter out variants with minor allele frequency < 1%, high missing genotyping rate (--geno 0.05), high missing individual rate (--mind 0.05) or variants that significantly deviated from the Hardy-Weinberg Equilibrium (--hwe 1e-06). Then, we also identified second-degree relatives (--genome, PI_HAT > 0.2) using a pruned set of variants in linkage disequilibrium (LD) (--indep-pairwise 100 10 0.1), and filtered out, from each pair, the individual with the lowest genotyping rate. Finally, we performed a Principal Components Analysis (PCA) of a pruned set of common variants of our study samples projected on the 1KGP Phase 3 samples on PLINK (version 1.9) (76,77), and removed individual outliers that did not cluster within the European sample of the 1KGP by inspecting the first three principal components (total PCA outliers across genotyping arrays = 81). Amongst the outliers, 63 individuals are from Quebec, 8 from British Columbia, 5 from the Atlantic Provinces, 5 from Alberta and none from Ontario.

### Imputation of Genotypes

Each genotyping array was first phased with EAGLE2 (version 2.0.5) (78) using the Sanger Imputation Server (79). After phasing, samples on each genotyping array were imputed on the Sanger Imputation Server using the positional Burrows-Wheeler transform (PBWT) algorithm (80) and the Haplotype Reference Consortium (HRC) release 1.1 dataset as reference (79). The HRC includes ~64,000 haplotypes and ~40,000,000 autosomal SNPs of ~32,000 individuals predominantly of European ancestry, which makes it ideal for the imputation of our datasets, which are of European-related ancestry. After imputation, we used PLINK (version 2) (76,77) to filter out variants with minor allele frequency < 1%, high missing genotyping rate (--geno 0.05), imputation score (INFO) < 0.3, or variants that significantly deviated from the Hardy-Weinberg Equilibrium (--hwe 1e-06).

### Phenotyping

Participants of the CanPath answered a questionnaire that included self-report on eye colour using the following discrete categories: grey, blue, green, amber, hazel or brown eye colour. These categories were then transformed into a linear scale using R (version 3.5.1) (81) to build a linear model with the following levels: 1 = grey or blue, 2 = green, 3 = hazel, 4 = brown. Supplementary Table 1 shows the number of individuals on each eye colour category by genotyping array. In addition, participants also reported their age and sex. We excluded the individuals who reported amber eye colour, due to the low sample count.

### Genome-Wide Association Studies (GWAS) and Meta-Analyses

Genome-wide association studies of eye colour were performed for each genotyping array with a linear mixed model on Genome-Wide Complex Trait Analysis (GCTA-MLMA) 1.26.0 (21,22), using an additive genetic model (i.e. the effect size is a linear function of the number of effect alleles). We performed a PCA of a pruned set of genotyped variants for each genotyping array after quality control, keeping only SNPs with MAF > 0.05 and excluding regions of high LD, using PLINK (version 1.9) (76,77). We included in the model sex, age and the first ten PCs as fixed effects, and a genetic relationship matrix (GRM) of genotyped SNPs computed on GCTA 1.26.0 (21,22) as random effects, to control for more subtle population structure. To evaluate the case of residual population substructure, we computed the expected vs. observed p-values using Q-Q plots on R (version 3.5.1) (R Core Team, 2019), and ran LD Score regression with LDSC, in which an LD Score intercept considerably higher than 1 may indicate remaining confounding bias (82).

We performed a meta-analysis of eye colour using the beta coefficient and standard error (SE) of each study on the software METASOFT (version 2.0.1) (23). METASOFT conducts a meta-analysis using a fixed effects model (FE), which works well when there is no evidence of heterogeneity (i.e. assumes same effect size across studies), and an optimized random effects model (RE2), which works well when there is evidence of heterogeneity among studies (23). Additionally, METASOFT computes two estimates of statistical heterogeneity, Cochran’s Q statistic and I^2^, as well as a Bayesian posterior probability that an effect exists on each individual study (M) (83).

For the meta-analyses results, we generated Manhattan and Q-Q plots using the qqman (84) and ggplot2 (85) R packages. In addition, we visualized the significant loci with regional plots using the web-based program LocusZoom (86), with the 1KGP Phase 3 European sample as reference LD. We focused our results on the fixed effects model, but we also report the RE2 on the summary statistics of the top signals as Supplementary File 1, and compared the statistical significance between both models when there was evidence of heterogeneity based on Cochran’s Q p-value and I^2^ statistics.

### Annotation of significant loci

We used the web-based program SNPNexus (87,88) to annotate the genome-wide significant signals (p-value < 1e-08) from the meta-analysis. Specifically, gene and variant type annotation were done using the University of California Santa Cruz (UCSC) and Ensembl databases (human genome version hg19); assessment of the predictive effect of non-synonymous coding variants on protein function was done with SIFT and PolyPhen scores. Both SIFT and PolyPhen output qualitative prediction scores (i.e. probably damaging/deleterious, possibly damaging/deleterious-low confidence, tolerated/benign). Non-coding variation scoring was assessed using CADD score, which is based on ranking the deleteriousness of a variant relative to all possible substitutions of the human genome. For instance, a score ≥ 20 indicates that the variant is predicted to be in the top 1% most deleterious variants in the genome (88). In addition, we explored the effect of significant loci on RNA and protein expression using the GTEx database (19) and the effect of significant genes using the Protein Atlas (89).

### Approximate Conditional Analyses of Association

In order to identify if the genome-wide significant loci of our original logistic meta-analyses were driven by one or more independent variants, we conducted approximate conditional and joint analyses of association (COJO) with GCTA (24). We performed the analysis (--cojo-slct) using as input the summary statistics of our eye colour meta-analysis (fixed effects, FE) and the weighted average effect allele frequency from all studies. In addition, the program requires a reference sample for computing LD correlations and, in the case of a meta-analysis, it is suggested to use one of the study’s large samples (24). Therefore, we ran the analysis twice: 1) using as a reference the sample genotyped with the Axiom UKBB array, and 2) using as a reference the sample genotyped with the GSA 24v1+MDP. We assumed that variants farther than 10 Mb are in complete linkage equilibrium and used a p-value threshold of 5e-08.

### Statistical Fine-Mapping of Significant Loci

We used the program FINEMAP (version 1.4) (90) to identify candidate causal variants in the genome-wide associated loci across the genome for eye colour. FINEMAP is based on a Bayesian framework, which uses summary statistics and LD correlations among variants to compute the posterior probabilities of causal variants, with a shotgun stochastic search algorithm (90). Compared to other methods, FINEMAP allows a maximum of 20 causal variants per locus. To run the program, we used as input the meta-analysis summary statistics, including the weighted average MAF across all studies, and an LD correlation matrix from one of the large samples in our study (Axiom UKBB array, N= 4,745). The LD correlation matrix was computed using LDStore (version 2.0), which considers genotype probabilities (32). We defined regions for fine-mapping as ± 500 kb regions flanking the lead SNP, based on the genome-wide and suggestive signals of association from the meta-analyses, and setting the maximum number of causal SNPs to 10 for each locus (i.e. a maximum of 10 credible sets). A credible set is comprised of SNPs that cumulatively reach a probability of at least 95%. The SNPs within a credible set are referred to as candidate causal variants and each of them has a corresponding posterior inclusion probability (PIP).

We filtered FINEMAP results by removing candidate causal variants with a log_10_BF < 2 from each of the 95% credible sets, where a log_10_BF indicates considerable evidence of causality. We annotated the remaining SNPs using SNPnexus (88) to obtain information about the overlapping/nearest genes, overlapping regulatory elements and CADD scores. Annotation of gene expression on ENCODE, Roadmap Epigenomics and Ensembl Regulatory Build was restricted to melanocytes and fibroblasts, which are the relevant cell types involved in eye colour. Based on the combined evidence of fine-mapping and posterior annotation, we defined the candidate causal variants with strong evidence of causality (based on their log_10_BF and annotation) as the most likely candidate causal variants. We computed LD correlations between the candidate causal SNP(s) on each locus (i.e. configuration with highest posterior probability and *k* number of SNPs) and the other variants on each locus using LDStore (version 2.0) and plotted the Posterior Inclusion Probability (PIP) results on R (version 3.5.1) (R Core Team, 2019) using ggplot2 (85).

Given that there may be putative functional SNPs that we did not genotype or did not impute with high accuracy, we explored if markers in the same LD blocks of the credible sets have functional annotations using HaploReg (version 4) (30,31). We used as input the most likely candidate causal SNP on each credible set, the LD from the 1KGP European population with a threshold of r^2^ ≥ 0.8 and the core chromatin 15-state model, which is based on several histone marks associated with promoters, enhancers, insulators and heterochromatin.

### Colocalization with Expression and Methylation QTLs from Cultured Melanocytes

We conducted colocalization analyses of our GWAS meta-analyses results with gene expression and methylation *cis-*QTL data from primary cultures of foreskin melanocytes, isolated from foreskin of 106 newborn males (61,62). *cis*-QTLs were assessed for variants in the ± 1Mb region of each gene or CpG. We used the program hyprcoloc (38) to obtain the posterior probability of a variant being shared between the eye colour GWAS signals and the expression or methylation QTLs. We tested all the significant eQTL genes or meQTL probes within ± 250 kb regions flanking the most significant GWAS SNP on each of the genome-wide regions (p-value ≤ 5e-8) from the meta-analysis summary statistics (five different loci). We used as LD reference the matrix obtained from the CanPath’s Axiom UKBB Array (INFO score > 0.3), computed on PLINK (version 1.9; --r square) (76,77). We kept colocalized regions that reached a posterior probability ≥ 0.8, indicating high confidence of shared causality.

### Transcriptome-Wide Association Studies

We performed a transcriptome-wide association study (TWAS) by imputing the expression profile of the CanPath cohort using GWAS summary statistics and melanocyte RNA-seq expression data (61). Using the program FUSION (39), we used as LD reference the CanPath’s Axiom UKBB genotyping array computed in binary PLINK format (version 1.9; --make-bed) (76,77). As recommended by FUSION, we used the LDSC *munge_sumstats.py* script to check the GWAS summary statistics (91). Before running the script, we filtered out SNPs with MAF < 0.01, SNPs with a genotyping missing rate > 0.01 and SNPs that failed Hardy-Weinberg test at significance threshold of 1 x 10^-7^ using PLINK (version 1.9; --maf 0.01, --geno 0.01, --hwe 10e-7) (76,77). We computed functional weights from our melanocyte RNA-seq data one gene at a time. Genes that failed quality control during a heritability check (using minimum heritability P-value of 0.01) were excluded from the further analyses, yielding a total of 3998 genes. We restricted the locus to 500 kb on either side of the gene boundary. We applied a significance cut-off to the final TWAS result of 1.25e-5 (i.e. 0.05/3998 genes tested). Finally, we performed conditional analysis on FUSION (FUSION_post.process.R script) if more than one gene in a locus was significant, to identify if these were independent signals.

### Genetic Correlations

We used a bivariate restricted maximum likelihood (REML) approach to test for genome-wide pleiotropy between hair and eye colour using GCTA (--reml-bivar option) (24). To consider the whole spectrum of colour in both traits, and thus maximize the number of loci, we coded both traits on a linear scale (excluding red hair colour). Hair colour ranged from 1= blonde, 2=light brown, 3= dark brown, and 4= black, whereas eye colour categories ranged from 1= grey/blue, 2= green, 3= hazel, and 4= brown. Given that the program requires genotype-level data, we computed the analysis twice, using the two largest samples for which eye colour data was available: Axiom UKBB array (N= 3,212) and GSA 24v1+MDP (N=2,429). Significance of the genetic correlations was computed with a likelihood ratio test on R (version 3.5.1) (81).

We first included in the model sex, age and the significant PCs as covariates, and restricted the analysis to SNPs with high INFO score (i.e. INFO > 0.8) and MAF > 1%. We then explored the correlations between each phenotype (i.e. hair and eye colour) and the eigenvectors of the principal components analysis. If the eigenvectors are correlated with the ancestry (i.e. geography) of the individuals, setting them as covariates may hinder the true genetic correlation between both traits, given that hair and eye colour are themselves correlated with ancestry. Therefore, we ran the genetic correlation a second time using as covariates only the non-significant principal components. In the case of the Axiom UKBB array we used PC3 and PC5, and in the case of GSA 24v1+MDP we used PC4, PC5 and PC8.

## Data Availability

We provide the genome-wide (p ≤ 5e-8) and suggestive (p ≤ 1e-6) signals identified in the eye colour metaanalysis as a Supplementary Information File (Supplementary File 1). Further information and requests for data published here should be directed to CanPath, which regulates the access to the data and biological materials (https://canpath.ca/). Melanocyte genotype data, RNA-seq expression data, and all meQTL association results are deposited in Genotypes and Phenotypes (dbGaP) under accession dbGaP: phs001500.v1.p1.

## Acknowledgements

The data used in this research were made available by CanPath – Canadian Partnership for Tomorrow’s Health (formerly CPTP), CARTaGENE, Alberta’s Tomorrow Project, Ontario Health Study, BC Generations Project and Atlantic PATH. The authors would like to thank all the participants of the Canadian Partnership for Tomorrow’s Health.

## Author Contributions

EJP and FLD designed the study. FLD, RT and EPC performed statistical analyses. FLD wrote the draft of the manuscript. FLD, EJP, EPC, KF, TZ, MAK, JC, IJ and KMB aided in the interpretation of the results and in the preparation of the final version of the manuscript.

## Financial Disclosure

FLD was supported by the National Council for Science and Technology (CONACYT) in Mexico. EJP received funding from the Natural Sciences and Engineering Research Council of Canada (NSERC Discovery Grant). RT, KF, MAK, JC, TZ, and KMB are supported by the Intramural Research Program of the NIH, National Cancer Institute, Division of Cancer Epidemiology and Genetics; https://dceg.cancer.gov/); the content of this publication does not necessarily reflect the views or policies of the Department of Health and Human Services, nor does mention of trade names, commercial products, or organizations imply endorsement by the U.S. Government. Computations were performed on the GPC supercomputer at the SciNet HPC Consortium, Canada and at the UTM High Performance Computing server at Mississauga, ON, Canada. This work also utilized the computational resources of the NIH HPC Biowulf cluster (http://hpc.nih.gov). SciNet is funded by: the Canada Foundation for Innovation under the auspices of Compute Canada; the Government of Ontario; Ontario Research Fund-Research Excellence; and the University of Toronto. The funders had no role in study design, data collection and analysis, decision to publish, or preparation of the manuscript.

## Supplementary Information

Supplementary Files, Tables and Figures are available for this paper.

## Notes

### Competing Interest Statement

The authors have declared no competing interest.

